# The first set of universal nuclear protein-coding loci markers for avian phylogenetic and population genetic studies

**DOI:** 10.1101/272732

**Authors:** Yang Liu, Simin Liu, Chia-Fen Yeh, Nan Zhang, Guoling Chen, Pinjia Que, Lu Dong, Shou-hsien Li

## Abstract

Multiple nuclear markers provide genetic polymorphism data for molecular systematics and population genetic studies. They are especially required for the coalescent-based analyses that can be used to accurately estimate species trees and infer population demographic histories. However, in avian evolutionary studies, these powerful coalescent-based methods are hindered by the lack of a sufficient number of markers. In this study, we designed PCR primers to amplify 136 nuclear protein-coding loci (NPCLs) by scanning the published Red Junglefowl (*Gallus gallus*) and Zebra Finch (*Taeniopygia guttata*) genomes. To test their utility, we amplified these loci in 41 bird species representing 23 Aves orders. The sixty-three best-performing NPCLs, based on high PCR success rates, were selected which had various mutation rates and were evenly distributed across 17 avian autosomal chromosomes and the Z chromosome. To test phylogenetic resolving power of these markers, we conducted a Neoavian phylogenies analysis using 63 concatenated NPCL markers derived from 48 whole genomes of birds. The resulting phylogenetic topology, to a large extent, is congruence with results resolved by previous whole genome data. To test the level of intraspecific polymorphism in these makers, we examined the genetic diversity in four populations of the Kentish Plover (*Charadrius alexandrinus*) at 17 of NPCL markers chosen at random. Our results showed that these NPCL markers exhibited a level of polymorphism comparable with mitochondrial loci. Therefore, this set of pan-avian nuclear protein-coding loci has great potential to facilitate studies in avian phylogenetics and population genetics.

---

Although the next generation sequencing technologies have produced sequences data in the unprecedented quantity with relative low cost^1^, traditional Sanger sequencing still has its niche in molecular evolutionary studies: pilot or small scale phylogenetic studies using PCR-based approach are cost-effectively and nearly available for every laboratory, beneficial to design sampling strategy and built an analysis scheme. By comparing molecular phylogenies based on different sizes of dataset, Rokas et al. ^2^ proposed that concatenation of a sufficient number of unlinked genes (>20) can overwhelm incongruent branches of the Tree of Life (TOL). Furthermore, tracing backwards from multiple genetic polymorphisms to find the most recent common ancestor (MRCA) of a group of individuals provides a sophisticated approach to clarify phylogenetic relationships among species (species tree approach) and to reconstruct the demographic history of populations^3,4^. However, the major drawback of this approach is the PCR performance of primers developed from one species is often unpredictable in the distantly related species; consequently, it is a time and cost consuming process to evaluate the performance of primers in a previously untested species. Therefore, a set of universal nuclear markers could provide an efficient way to ease this time consuming process. It should greatly facilitate the use of coalescent-based analyses to answer phylogenetic and population genetic questions^5^.

Nuclear Protein-coding Loci (NPCLs) are exons without flanking introns^6^, and are widely used in interspecific phylogenetic studies (e.g. *RAG1*^7^, *c-myc* ^8,9^). NPCL markers possess favorable properties including homogeneous base composition, varied evolutionary rates and easy alignment across species or populations^10,11^. Moreover, orthologous genes can be identified accurately using their annotations ^12,13^. Several sets of universal NPCL markers had been developed specially for beetles^14^, fish^15^, reptiles^6^, amphibian and vertebrates^16,17^. However, there is still no sufficient number of easily amplifiable NPCL markers that can fulfill the needs of modern coalescent-based analysis for most of bird species. As the most common and species-rich group of terrestrial vertebrates, birds exhibit tremendous diversity in their phenotypes, ecology, habitats and behaviors^18^. So far, a considerable effort has been devoted to resolve the phylogenetic relationships from higher taxonomic categories^19^-^21^to sister species^22-26^. In addition to phylogenetics, modeling-based approaches using multiple nuclear genes have also shed light on population structure and demographic history and allowed inferences of selection pressures in non-model organisms^27-30^. The rapid advance in these sub-disciplines in evolutionary biology always hinges upon proper sampling design and a rigorous statistical approach, but it also requires data on multiple independent loci with an appropriate level of genetic polymorphism^31^, which allows the application of sophisticated modeling and thus hypothesis testing.

Efforts of developing universal PCR primers have facilitated avian phylogenetic and population genetic studies^32-34^. For example, Dawson et al. ^35^ developed a set of microsatellite markers with high cross-species utility, suitable for paternity and population studies. Backström et al. ^36^ developed more than 200 exons flanking introns, which were evenly distributed throughout the avian genome.

However, a variable number of indels (insertions and deletions) in the intron complicate the subsequent amplification, sequencing and alignment of these exons. Conserved and easily aligned exonic regions are ideal alternatives to compensate for resolving power for phylogenetic reconstruction^13^. Kimball et al. ^37^tested the utility of 36 published markers on 42-199 bird species with only five exonic markers therein. Kerr et al. ^38^developed 100 exonic markers from five avian genomes, and finally tested a subset of 25 markers in 12 avian orders. The quantity of NPCL markers is far from adequate as exon length should be longer than intron sequences to yield sufficient phylogenetic resolution^39^. Using a small number of universal NPCL markers could increase the probability of error when estimating species relationships due to the conflict of gene tree topologies. To overcome the problem, it has been advocated to use more genes with longer sequences^40^. However, some obstacles have hindered the development of universal NPCL markers. Firstly, widespread flanking introns make the identification of the exon boundaries of a specific NPCL marker difficult^6^. Secondly, multiple nuclear loci are required to be distributed evenly and widely across the whole genome in order to indicate a variety of historical signals. And finally, low-cost and easy amplification are important requisites. The development of a universal set of NPCL markers for birds should significantly reduce the time required for future research as well as its cost, and facilitate the application of coalescent-based methods in avian evolutionary studies.

In this study, we aimed to develop a set of avian universal NPCL markers that can be widely utilized in avian phylogenetic and population genetic studies. By comparing the published genomes of the Red Junglefowl (*Gallus gallus*) and the Zebra Finch (*Taeniopygia guttata*), we designed 136 pairs of NPCL primers and amplified them in 41 species representing 23 avian orders to check their versatility. To test the resolving power of these markers, we further constructed a phylogenetic tree and estimated mutation rates by extracting universal NPCLs from 48 published avian genomes^41^. Moreover, samples from four populations of the Kentish Plover (*Charadrius alexandrinus*) were also amplified to estimate the intra-specific polymorphic level of these universal NPCLs.

## Results

### Pan-avian order amplifications of the novel NPCLs

The genome alignment and BLAST procedures resulted in 136 NPCL candidates, which were broadly distributed across 24 autosomal chromosomes and the Z chromosome of the Zebra Finch genome. Their original fragment length ranged from 815bp to 7176bp (Supplementary Table S1). We thus nominated each NPCL marker using abbreviation of the associated protein-coding regions according to gene annotation of Zebra Finch (Supplementary Table S1). More than one primer pairs were conducted for each NPCL marker candidate, and we finally chose the pair of PCR markers with the highest score denoting the level of conservatism between Zebra Finch and Red Junglefowl genomes.

In total, 5,146 PCRs were performed to amplify the 136 NPCLs in 41 species representing 23 avian orders (Fig. 1A). Among them, 2,875 (55.9%) of PCR performances produced a target band (Supplementary Table S3). For the 136 candidates, we successfully amplified 12 NPCLs in all 23 orders, with 100% PCR success rate (PSR). Sixty three of the 136 candidate NPCL markers had a relatively good overall PCR performance (PSR ≥ 80%) (Fig. 2A); all of them were successfully amplified in *Caprimulgiformes* and *Gruiformes*, and the PSR ranged from 65% to 97% in other orders (Fig. 1B, Supplementary Table S3). This set of 63 universal avian nuclear markers was distributed across 17 autosomal chromosomes and the Z chromosome (Fig. 3).

**Figure 1.**
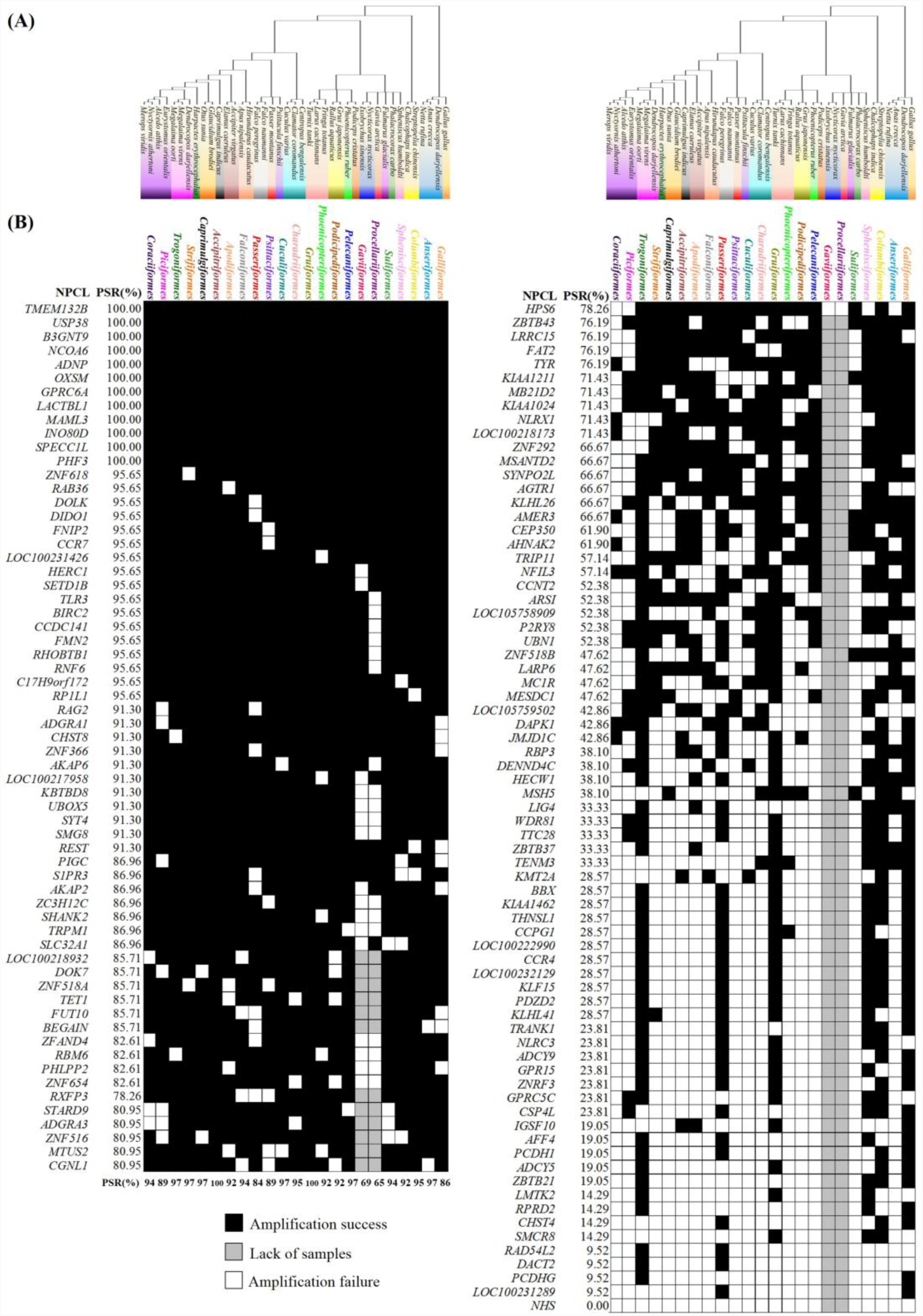
PCR performance for the 136 NPCL marker candidates in 23 avian orders. **(A)** Genetic relationships among our experimental samples. 41 species are highlighted in different colors representing 23 avian orders widely distributed in the avian phylogenetic tree. **(B)** PCR performance for 136 NPCL marker candidates. Each square represents a PCR result. Success is shown in black and failure in white. 430 of 5146 reactions that could not be produced due to a paucity of DNA are shown in grey. The gene name and PCR success rate of each NPCL marker are indicated to the left. The success rate of each avian order is indicated at the bottom of the matrix of 63 universal NPCL markers.

**Figure 2.**
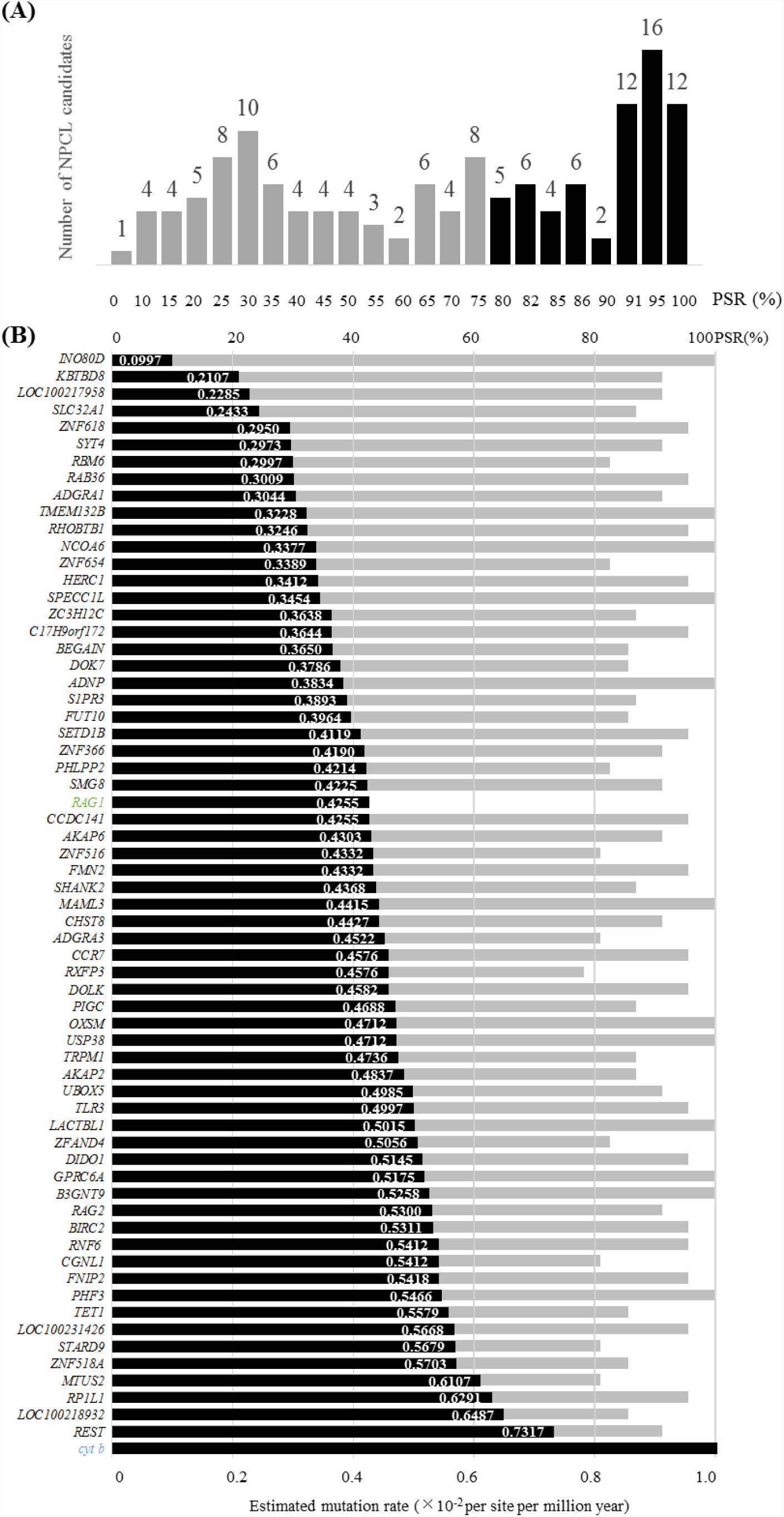
PCR success rate distribution for the 136 NPCL marker candidates and mutation rates for the 63 universal NPCL markers. **(A)** PCR success rate distribution for 136 candidates in 23 avian orders. The 63 NPCL markers with PCR success rates higher than 80% are shown in black; other loci with PSR success rates below 80% (in grey) were excluded from the subsequent analysis. The number above each bar shows the number of NPCL marker candidates. **(B)** Mutation and success rates for the 63 universal NPCL markers. The markers were sorted according to estimated mutation rates (bars in black) from low to high. The number on the right of each bar is the mutation rate of each NPCL marker. PCR success rates are shown underneath (bars in grey). The mutation rates of widely-used NPCL *RAG1* (in green) and mitochondrial gene *cyt b* (in blue) were selected as references.

**Figure 3.**
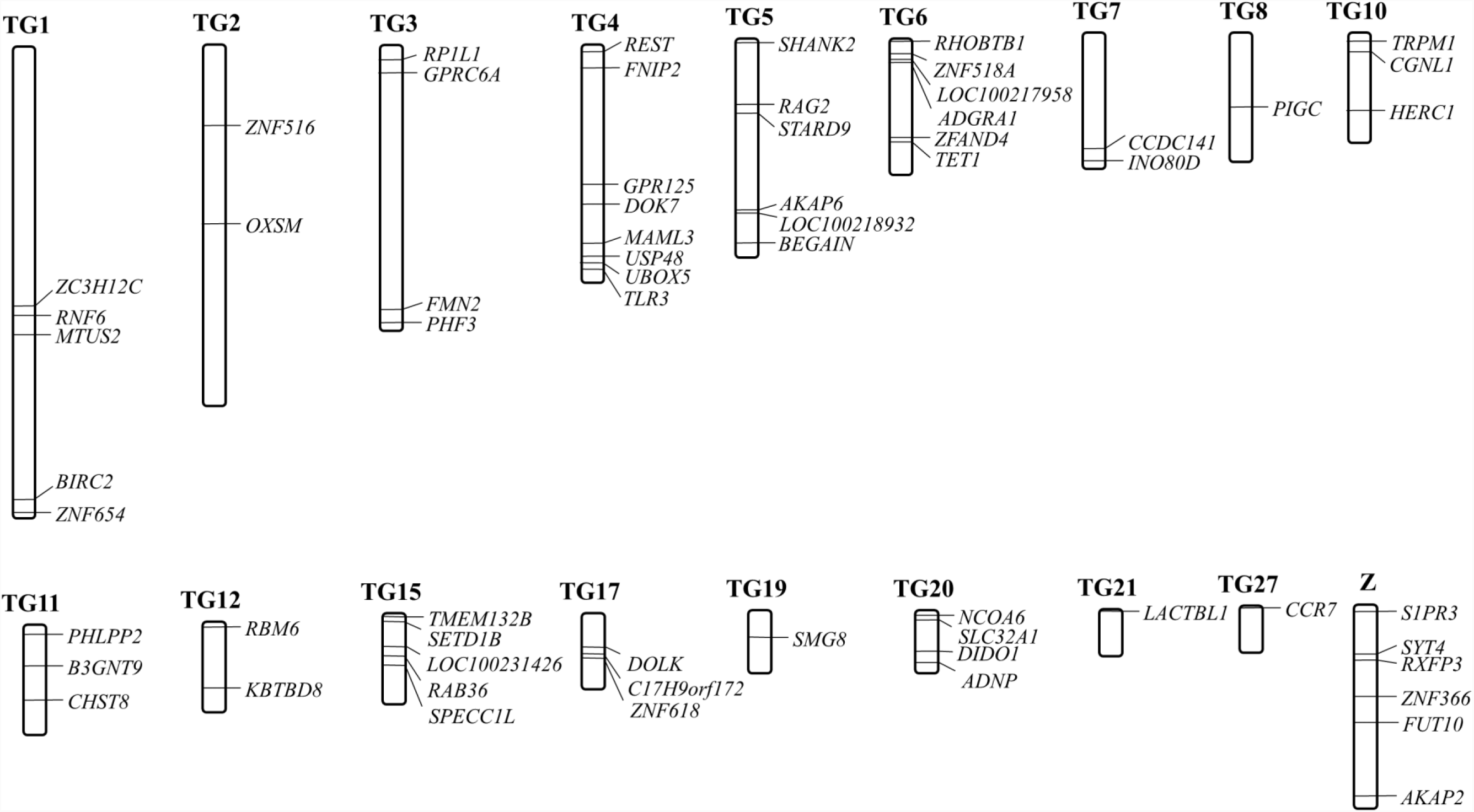
Chromosome mapping of the 63 avian universal NPCL markers in the genome of Zebra Finch (*Taeniopygia guttata*). The 63 universal NPCL markers with more than 80% PCR success rate were widely distributed in 17 autosomal chromosomes and the Z chromosome.

### Interspecific mutation rate and phylogenic construction of the 63 universal NPCLs

The genome-based BLAST results showed that the widely used genetic markers, cytochrome *b* (*cyt b*) of mitochondrial DNA(mtDNA) and *RAG1*, an extensively used nuclear gene42,43 were located in all 48 published genomes^41^. For the newly developed NPCLs, we located 56 loci across all 48 avian genomes. Among the remaining seven NPCLs, six of them were located in 47 genomes and two missing data recorded at the locus *FUT10*. Combined, BLAST results confirmed the set of 63 universal NPCL markers were orthologous among these 48 species (Supplementary Table S4) and the resulting concatenated matrix with sequences of approximately 96 kb was obtained.

The range of the estimated mutation rates for the universal avian NPCLs is broad; it ranged from 0.0997 to 0.7317 ×10 ^−8^ per site per million years (Fig. 2B). Among these 63 NPCLs, the mutation rates of 27 were slower than the mutation rate of *RAG1*, whilst the other 36 NPCLs were faster. All NPCLs showed a slower mutation rate than that of the mitochondrial *cyt b*.

We constructed a Maximum Likelihood (ML) tree based on 63 concatenated NPCLs from 48 species, representing 34 orders of extant birds (Fig. 4). The resulting topology is largely similar with the recent phylogenomic studies^41,44,45^. Neoaves and Galloanseres, which united in the infraclass Neognathae, as well as Palaeognathae were three major groups with highest bootstrap support (100%). Among Neoaves group, two major clades, core landbirds (Telluraves) and core waterbirds (Aequornithia) were strongly supported by whole-genome data^41^ and 259 independent nuclear loci^44^. Within core landbirds, the clade containing Passerimorphae (Passeriformes + parrots), Falconiformes (falcons), Cariamiformes (seriemas) is sister to Craciimorphae (bee-eaters + woodpeckers + hornbills + trogons + cuckoo-roller + mousebirds), which is paraphyletic to Strigiformes (owls) and its sister clade Accipitrimorphae (eagles + New World vultures). Within core waterbirds, Pelecanlmorphae (pelicans + herons + ibises+cormornts) and Procellariimorphae (fulmars + penguins) are two monophyletic groups, sister to Gaviimorphae (loons). Other clades such as Phoenicopterimorphae (flamingos + grebes), Otidimorphae (bustards + turacos + cuckoos), Caprimulgimorphae (hummingbirds + swifts + nightjars) and Phaethontimorphae (tropicbirds + sunbitterns) are identical with previous studies^41,45,46^. However, we also found discordances between this phylogenetic tree and previous results^41,44-46^, specifically in some branches with conflict placements with low support. For example, Columbiformes (doves), Pterocliformes (sandgrouses) and Mesitornithiformes (mesites) are not clustered into Columbimorphae. The placement of Charadriiformes (plovers), Gruiformes (cranes) and Opisthocomiformes (hoatzins) are incongruence with Jarvis et al^41^, respectively.

**Figure 4.**
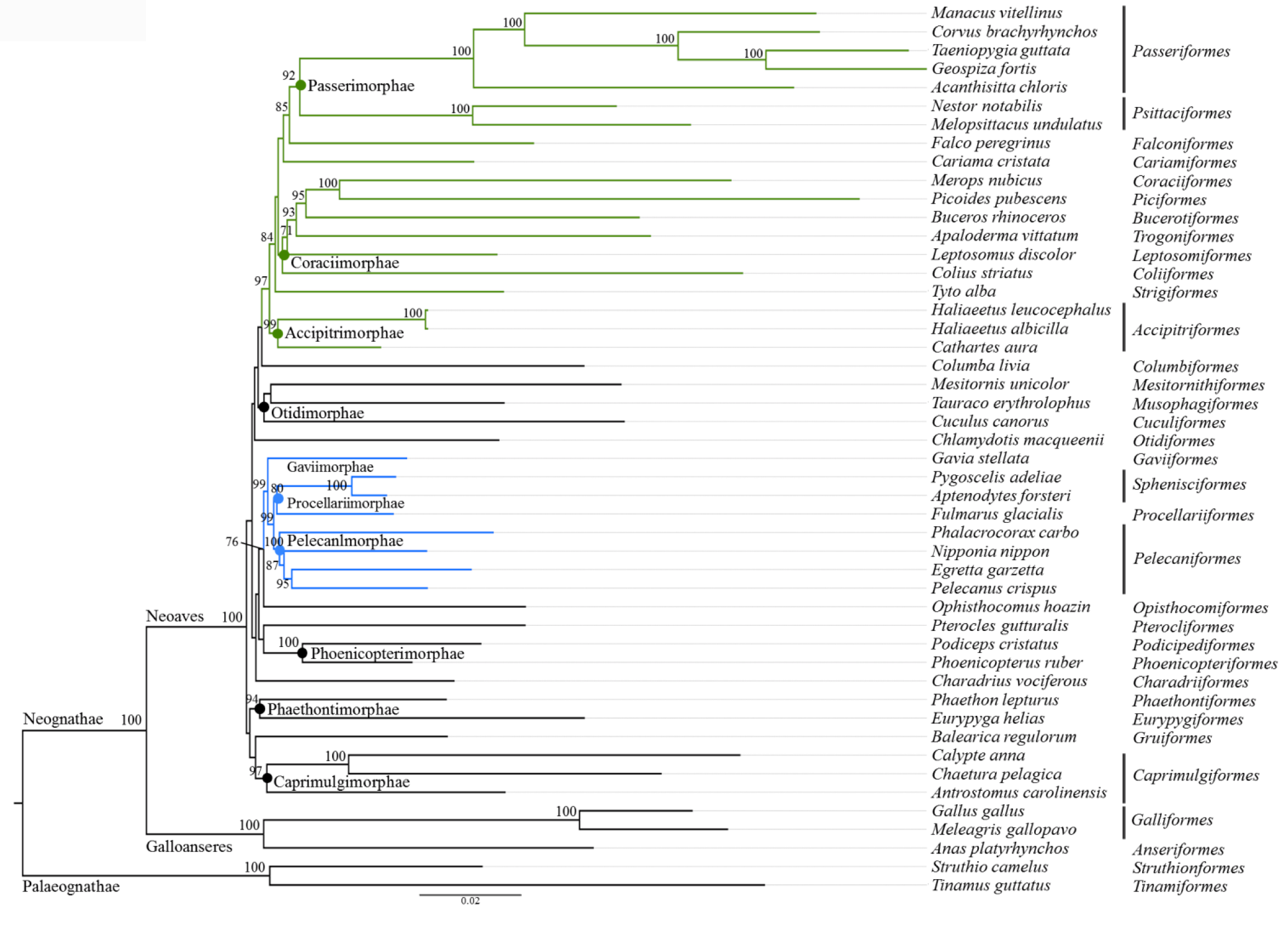
Phylogenetic analysis of Neoaves using 63 NPCLs from 48 bird species. The names of species are abbreviated (refer to supplementary Table S4), representing 30 orders of Neoavian birds. Superorders are labelled on the nodes as classification in previous study^41^ using genomic data. Bootstrap support over 70% are indicated above nodes.

### Intraspecific polymorphism of 17 randomly selected NPCL markers

A total of 12,420bp DNA sequences, including 11,196bp of 17 NPCLs and 1,224bp of two mitochondrial loci were sequenced in 40 samples representing four populations of the Kentish Plover. The NPCL markers showed varied degrees of polymorphism, with the exception of locus *KBTBD8* (Fig. 5). There were 10 polymorphic sites in loci *BIRC2* and *FMN2*, while there were only 1-6 polymorphic sites in other loci. Correspondingly, *BIRC2* and *FMN2* possessed the highest values of haplotype and nucleotide diversity (mean *Hd*= 0.84 and 0.92, mean *π*= 0.0047 and 0.0040, respectively). In contrast, a mitochondrial gene *ND3* had only one polymorphic site, yielding a low haplotype (mean *Hd*= 0.36) and nucleotide diversity (mean *π=* 0.0009). When we combined the results on interspecific mutation rates and intraspecific polymorphism, we found that the inter-and intraspecific genetic diversity of our gene markers were incongruent; although the estimated mutation rates at the study NPCLs were all much lower than that of mitochondrial gene *cyt b*, the intraspecific polymorphism at nine NPCL markers was higher than that of two mitochondrial genes.

**Figure 5.**
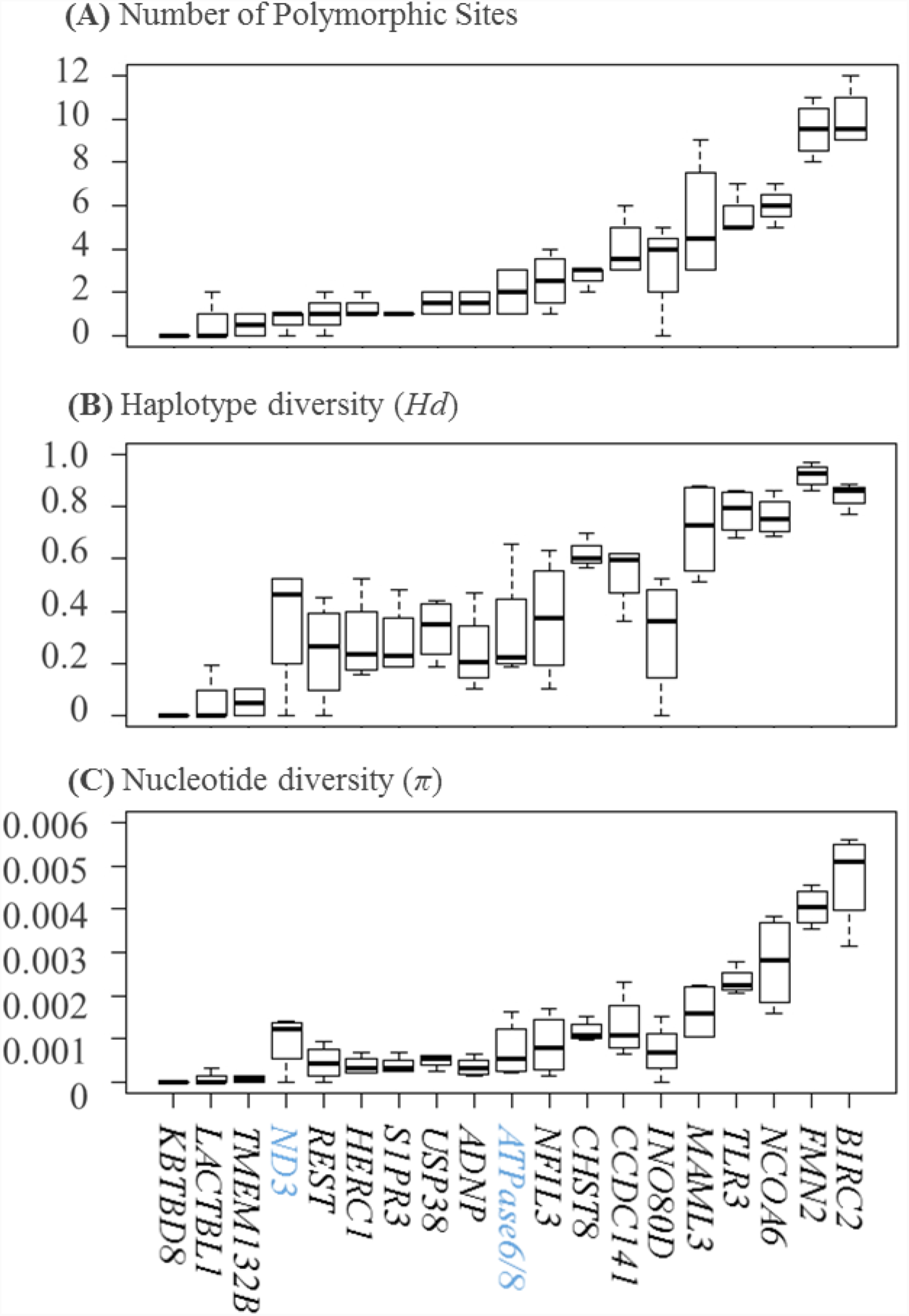
Polymorphism at 17 avian universal NPCL markers and two mitochondrial loci in four populations of Kentish Plover (*Charadrius alexandrinus*). Markers were sorted according to their level of polymorphisms, from low to high. Gene names can be found at the bottom of the three box plots. Mitochondrial genes (in blue) were selected as references **(A)** Average number of nucleotide sites ranging from 0 to 10. **(B)** Haplotype diversity ranging from 0 to 0.97. **(C)** Nucleotide diversity ranging from 0 to 5.59.

The genetic polymorphism parameters varied greatly, not only among genes, but among populations as well. For example, the *Hd* value of the *MAML3* gene was lowest in the Taiwan population (*Hd*= 0.51) and highest in the Qinghai population (*Hd*= 0.88). The measure for nucleotide diversity, *π* of the *NCOA6* gene was lowest in the Taiwan population (*π*= 1.59) and highest in the Guangxi population (*Sπ*= 3.81). Detailed information on measures from each population is available in Supplementary Table S5. The HKA test suggested no departure from the neutral expectation for any of the 17 NPCL markers. Similarly, the test of Tajima’s *D* showed that none of the 17 NPCL markers deviated significantly from neutrality (Supplementary Table S5).

## Discussion

We developed a set of 63 avian universal NPCL markers with diverse mutation rates and levels of intraspecific polymorphism. Our results showed that the 63 NPCL markers were successfully amplified in most of the species tested, representing 23 extant orders across major lineages of the avian tree of life (PCR success rate >80%), and denoted different levels of inter-and intraspecific polymorphism. Therefore, our NPCL set will provide a highly versatile genetic toolkit for a broad range of molecular phylogenetic and ecological applications. Moreover, the genetic marker system we provide here is cheap and easy to apply. Any molecular laboratories that are capable of performing PCRs can adopt our marker system effortlessly. Hence, this novel set of universal NPCL markers has great potential to be widely applied in evolutionary biology studies in birds.

Inherited from different chromosomes, concatenation of nuclear markers contribute multiple independent estimates to species trees^47,48^, in order to alleviate the node conflicts of gene trees caused by incomplete lineage sorting, horizontal gene transfer, inconsistent evolutionary rates, gene duplication and/or gene loss and so on^49,50^. We constructed an avian phylogenetic tree using concatenation of 63 NPCLs across 18 chromosomes from 48 genomes. The result is largely similar with previous phylogenomic works using different data types like multiple nuclear loci^41,44^, introns^20^, ultraconserved elements (UCEs) ^46,51^and retroposon presence/absence matrix^45^. The congruent parts of topology reveal multiple cluster clades, such as Telluraves (core landbirds), Aequornithes (core waterbirds) and Phoenicopterimorphae, Otidimorphae, Caprimulgimorphae and Phaethontimorphae. We also find some unresolved placements comparing with Jarvis et al^41^. These include Columbiformes (doves), Pterocliformes (sandgrouses), Mesitornithiformes (mesites), Charadriiformes (plovers) and Gruiformes (cranes), which exhibits hard polytomies in the avian tree of life. Though recent efforts in avian phylogenomic studies using whole-genome^41^ or genome-level data^44^, irresolvable relationships have been found in some clades^41,44-46^. Suh et al. ^45^ investigated the causes of phylogenetic irresolvabilities and concluded that such phylogenetic discordances were originated from prevalent ancestral polymorphism denoted by incomplete lineage sorting (ILS) ^52^, which is probably associated with an initial near-K-Pg super-radiation^41^ in Neoaves. Unlike the two other main radiations that gave rise to the core waterbirds and core landbirds clades, the massive near-K-Pg super-radiation in Neoaves, containing several unresolved lineages, leads extreme ILS and associated network-like phylogenetic relationships^45-46^. On one hand, again, the topology reconstructed by the present set of universal NPCL markers captures these patterns, and suggests hard polytomies due to biological limitation of phylogenetic methods. On the other hand, it implies that our NPCL markers have sufficient polymorphism to resolve phylogenetic relationships among lineages with less ILS in Neoaves.

We also found that this set of novel NPCL markers has the potential to be applied in population genetic studies, in which researchers usually prefer to use abundant markers with high mutation rates. For example, microsatellites were developed for specific species or orders to detect differences in genotypes and further to quantify intraspecific genetic diversity^35,53,54^But introns like microsatellites have high levels of length homoplasy^55^. It is commonly assumed that NPCLs are conservative loci, highly suitable to address questions concerning high-level systematics^40^. However, some population genetic studies highlight the importance of using functional exonic SNPs in population genetic studies^11,56^, comparing to neutral markers (such as microsatellite and mitochondrial DNA). Datasets that contain numbers of several to more than 100 exon genes^51,57^can support accurate and reliable estimates of population genetic parameters^55,58^, and have a substantial power in population genetic analysis^5,40,50,59^Studying 17 NPCL markers in the Kentish Plover, we found that sixteen had low to moderate levels of intraspecific polymorphism and nine of them showed higher genetic diversity than mitochondrial genes in this study. Although a previous study showed a low level of genetic differentiation across Eurasian populations^60^, this species exhibits variability in morphology and behavior among and within populations in East Asia^61^, warranting further coalescent-based analysis of their evolutionary history. This dataset provides sufficient information to study population genetics in the Kentish Plover in East Asia.

In order to infer correct phylogenetic relationships in different taxonomic levels, it is essential to choose unlinked genes with different mutation rates^62^. Our novel set of NPCL markers offer a wide range of mutation rates. The comparisons of mutation rates between the new NPCL markers with commonly used nuclear loci *RAG1*^7^ and some loci at mtDNA^63^ provide a reference for marker choice (Figure 2B, 5). In principal, it is advisable to use markers with slow mutation rates to resolve deep nodes and fast mutation rates to population genetic studies. Moreover, coalescent theory is widely used to estimate species tree and population demographic parameters, such as divergence times^64,65^and effective population sizes (*N*e) ^59^. The associated analyses, such as species-tree estimation, e.g. MP-EST^66^, *BEAST^67^, BP&P^68^, and demographic analysis, such as Isolation with Migration (IM) model^69,70^and Approximate Bayesian Computation (ABC) simulations^59^ require multiple independent loci with different demographic histories and mutation rates. In this regard, markers from different genomic segments, such as introns (developed previously^35,37^) and exons we developed are preferred to combine and to be used. It is no doubt that the present marker set is a useful resource to generate multilocus datasets for avian evolutionary studies in different taxonomic levels. In fact some studies have used these novel NPCL markers to apply the aforementioned analyses^25,30^.

Compared with traditional Sanger sequencing, the fast development of next generation sequencing (NGS) techniques has enabled researchers to obtain genetic polymorphisms easily^71^. For example, Jarvis et al^41^performed a highly resolved phylogenetic tree of 48 species using phylogenomic methods, and Prum et al. ^44^ conducted a comprehensive phylogeny of 198 species within the Neoaves, which diversified very quickly, using genome-scale data by targeted NGS. Multilocus methods do not use as much as genomic data. However, we consider that this set of universal NPCL markers has its niche in avian molecular studies. Sanger sequencing technique of NPCL markers is less sensitive to the quality of template DNA like sequence capture approach than other genomic approaches. Degraded DNA or a small quantity of DNA is also workable, like feather and museum samples. It is always a tradeoff between template DNA quality and PCR product length. With the novel NPCL markers, we aimed to amplify a fragment of 700-1200 bp sequence of each locus. Hence they should be applicable to avian blood, tissue and feathers. Moreover, a thorough analysis pipeline for traditional PCR-based method is available, supported by a series of visualized operating software, e.g. MEGA, DNASTAR, DnaSP, BEAST and etc., which are widely used in molecular phylogenetic analysis. Processing genomic data always places high demands on bioinformatics and computational power72. High-quality samples for NGS, project budget, bioinformatic facilities are not available to all laboratory. It is still useful and necessary to align orthologous sequences across multiple hierarchical levels using NPCL markers, especially for a pilot or small scale study.

However, there are some limitations when using this set of universal NPCL markers. Firstly, PCR performances were simultaneously tested under a unified protocol (e.g. Tm=50°C), so that the PSR of each NPCL marker might be underestimated. Reducing the annealing temperature by 1~2°C would improve the success rate in practice. There is also the possibility that PCR produced target sequences but also non-specific amplicons. We could slightly raise the annealing temperature to increase specificity or perform extra steps including gel purification and cloning. Furthermore, the interspecific polymorphic parameters of the 17 NPCL markers are reference values for Kentish plovers. Different evolutionary forces, such as genetic drift or natural selection, can act on different regions of the genome, causing a various evolutionary rates and demographic histories in different species^73^.

Thus, different combinations of markers are important for specific questions. For example, NPCL markers on the Z chromosome could be selected to solve questions involving sexual selection and mate choice. There is also a trade-off between the number of markers and time-and cost-efficiency. In avian phylogenetic analysis, random errors can be reduced by employing more markers, whilst, as a consequence of this procedure, systemic errors would increase due to differences in nucleotide composition and various mutation rates^74^. Kimball et al. proposed that adopting various analytical methods might overcome these adverse effects^75^.

In conclusion, we have developed 63 avian universal NPCL markers, evenly distributed across 17 autosome chromosomes and the Z chromosome. This set of universal NPCL markers had high PCR success rates (PSR>80%) in 23 avian orders. Its wide range of mutation rates are suitable to resolve phylogenetic relationships at both low and high-level. Furthermore, various intraspecific polymorphisms are potentially useful to provide deep-level divergence and demographic information for population genetics. Though high-throughput genetic polymorphism data from next generation sequencing undoubtedly provide a more comprehensive vision for avian evolutionary history and genomic patterns, we believe that this set of exonic markers provides a relatively reliable and repeatable solution and could have widespread application in phylogenetic and population genetics studies.

## Methods

### Development of NPCL markers and primer design

To screen NPCL marker candidates, we aligned parts of the genome of two species with a distant phylogenetic relationship^76^, the Red Junglefowl (GCA_000002315) and the Zebra Finch (GCA_000151805). Firstly we identified long (>600bp) single-copy exons within the genome of the Zebra Finch and took these exons as templates. Then we aligned them with the genome of the Red Junglefowl using BLAST (Basic Local Alignment Search Tool). We assumed that query sequences of the Red Junglefowl with identity more than 80% and length more than 50% of templates length were orthologous exons and employed them as NPCL marker candidates.

We used the program Primer3^77^ to design the primers for NPCL marker candidates. We selected exon sequences of Zebra Finch as templates, focusing on High-scoring Segment Pairs region (700-1200bp). For each primer pair, the oligomer ranged from 18bp to 25bp and GC content ranged from 20% to 80%. Furthermore, we tested a single primer for self-complementarity by setting complementarity score to less than 6.00, so as to predict the tendency of primers to anneal to each other without necessarily causing self-priming in the PCR. Complementarity 3’ score was set as default (< 3.00) to test the complementarity between left and right primers.

### Tests on the universality of the NPCL markers

To test the amplification performance of these new NPCL markers, we selected 41 species of 23 representative Aves orders (Supplementary Table S2). We used a set of 10000 trees with 9993 operational taxonomic unites (OTUs) downloaded from http://birdtree.org/ to demonstrate the phylogenetic relationships among selected species^18^.

Total genomic DNA was extracted from ethanol-preserved muscle tissue or blood using a TIANamp Genomic DNA kit (TIANGEN, China) and stored at 4?. DNA concentration and purity were estimated by NanoDrop 2000 (Thermo Scientific, USA). We used Touchdown PCR (TD-PCR) ^78^, an improved standard PCR to test the utility of primers sensitively, by decreasing the annealing temperature 1°C /cycle from Tm+10°C to Tm (Melting Temperature). The Touchdown PCR was performed in a Veriti96 PCR thermal cycler system (ABI, USA) using a 10μl reaction containing 2μl template DNA (10-70ng totally), with mixed concentrations of 10×PCR buffer, 20μM dNTP, 10mM of each forward and reverse primer, and 5U Taq polymerase (Takara, China). The initial temperature profile was 2 min at 94°C, 10 cycles at 94°C for 30s, 60-50°C (decreasing the annealing temperature by 1°C per cycle) for 30s and 72°C for 90s followed by 30 similar cycles but with a constant temperature of 50°C. This process was concluded with an extra elongation step at 72°C for 10 min. A successful amplification was recorded if a single clear band (target locus) was observable under ultraviolet light after being isolated on a 1% TAE agarose gel at 120V for 30 min.

### Estimation of inter-species mutation rates and Construction of Neoavian phylogeny

We downloaded 48 avian genomes covering all orders of Neoaves^41^. Sixty-five gene sequences, including the 63 universal NPCLs and two frequently-used genes (*RAG1*, and mitochondrial cytochrome *b* (*cyt b*)) as control, were retrieved and aligned in these genomes against genes in Zebra Finch (abbreviation as *Tgu1*) using BLAST. These sequences were extracted and filtered in batches by own-developed Perl script in the Tianhe-2 server (School of Advanced Computing, Sun Yat-sen University). Because of different genome sequence format, the script was unable to retrieve sequences in four species, i.e. *Anas platyrhynchos, Gallus gallus, Meleagris gallopavo* and *Melopsittacus undulates*^41^. Hence, we manually searched and obtained orthologues of the four species using BLAST tool on the website https://blast.ncbi.nlm.nih.gov/Blast.cgi. We further aligned the obtained NPCL orthologous sequences using MEGA v6.0^79^.

To estimate the mutation rate of each gene, we firstly computed the overall mean genetic distance at each NPCL marker in MEGA v6.0 with 1000 bootstrap replicates. Then we calculated the ratio of genetic distance between each NPCL and *cyt b*. Finally, we multiplied the ratio by the average mutation rate in the *cyt b* (0.01035 mutations per site per million years) ^80^ to get the average mutation rate of each NPCL^81^.

To construct the phylogenetic relationship of Neoavian birds in order level, we concatenated all NPCL sequences and reconstructed the maximum likelihood tree by RAxML v8.2.1^82^, with GTRCAT model and 1,000 bootstrap runs. Maximum-likelihood-bootstrap proportions ≥ 70% were considered strong support^83^.

### Intra-species polymorphism measurements

We amplified 17 random NPCLs in 40 Kentish Plover (*Charadrius alexandrinus*) blood samples from live-trapped birds in a noninvasive manner. To compare our data with previous genetic analyses on European populations of the Kentish Plover^84^ and compare the degree of polymorphism between nuclear and mtDNA, we added two mtDNA loci, ATPase subunit six concatenated with partial ATPase subunit 8 (*ATPase6/8*) and NADH dehydrogenase subunit 3 (*ND3*). Blood samples were collected from four breeding populations of plovers, Guangxi (GX), Qinghai (QH), Hebei (HB) and Taiwan (TW) (Table S2). The same protocol for DNA extraction and PCR amplification was followed as above, and the products were sequenced on ABI3730XL (Applied Biosystems, USA) by Beijing Genomics Institute (BGI, China).

Both strands of the amplicons were assembled, and the heterozygosity of nuclear genes was detected using SeqMan v7.1.0.44^85^. Some parameters of the DNA polymorphism, the number of polymorphic sites (*S*) and haplotypes (*H*), haplotype diversity (*Hd*), and nucleotide diversity *(π)* were calculated using DnaSP v5.0^86^. The neutrality of each locus was tested using the Hudson-Kreitman-Aguade (HKA) test^87^ and Tajima’s *D*^88^ implemented in DnaSP v5.0.

## Acknowledgements

We are grateful to Per Alström, Fuming Lei, Fasheng Zou, Xiaojun Yang, Ulf Johansson, Chung-Yu Chiang, Jonathan Reeves, Yingyong Wang, Menxiu Tong, Qin Huang, Zhechun Zhang, Xuejing Wang, Xin Lin, Jian Zhao for supplying tissue, blood or DNA samples used in this study, and Zhenhao Luo for providing technical support to BLAST script methods, and Alan Watson for editing the text. This study was supported by the National Science Foundation of China (No. 31301875 & No. 31572251 to Yang Liu; No. 31471987 to Lu Dong, No. 31600297 to Pinjia Que), and the National Key Program of Research and Development, Ministry of Science and Technology Grant 2016YFC0503200 to Lu Dong, and some DNA samples of birds were collected during ‘The Comprehensive Scientific Survey of Biodiversity from Luoxiao Range Region in China (2013FY111500)’. Computational work was funded by Special Program for Applied Research on Super Computation of the NSFC-Guangdong Joint Fund (the second phase) under Grant No. U1501501 to Yang Liu.

## Author contributions statement

Y.L., SH.L. and D.L. designed this study. CF. Y. and N.Z. carried out the primer design and BLAST procedures. GL.C. and PJ.Q. provided materials and technical support in the lab. SM.L. completed wet lab experiments, analyzed the data and wrote the manuscript with Y.L.

